# A macrophage response to *Mycobacterium leprae* phenolic glycolipid initiates nerve damage in leprosy

**DOI:** 10.1101/127944

**Authors:** Cressida A. Madigan, C.J. Cambier, Kindra M. Kelly-Scumpia, Philip O. Scumpia, Tan-Yun Cheng, Joseph Zailaa, Barry R. Bloom, D. Branch Moody, Steven T. Smale, Alvaro Sagasti, Robert L. Modlin, Lalita Ramakrishnan

**Affiliations:** Division of Dermatology, Department of Medicine, David Geffen School of Medicine, University of California, Los Angeles, CA, USA, 90095.; Department of Microbiology, Immunology and Molecular Genetics, David Geffen School of Medicine, University of California, Los Angeles, CA, USA, 90095.; Department of Microbiology, University of Washington, Seattle, WA 98195, USA.; Department of Immunology, University of Washington, Seattle, WA 98195, USA.; Division of Rheumatology, Immunology and Allergy, Brigham and Women’s Hospital, Harvard Medical School, Boston, MA, USA.; Harvard School of Public Health, Boston, MA 02115, USA.; Department of Molecular, Cell and Developmental Biology, University of California, Los Angeles, CA 90095, USA.; Department of Medicine, University of Washington, Seattle, WA 98195, USA.; Molecular Immunity Unit, Department of Medicine, University of Cambridge, MRC Laboratory of Molecular Biology. Cambridge UK CB2 OQH, UK.; Molecular Biology Institute, University of California, Los Angeles, CA, USA, 90095.

## Abstract

*Mycobacterium leprae* causes leprosy, and is unique among mycobacterial diseases in producing peripheral neuropathy. This debilitating morbidity is attributed to axon demyelination resulting from direct interactions of the *M. leprae*-specific phenolic glycolipid 1 (PGL-1) with myelinating glia, and their subsequent infection. Here, we use transparent zebrafish larvae to visualize the earliest events of *M. leprae*-induced nerve damage. We find that demyelination and axonal damage are not directly initiated by *M. leprae* but by infected macrophages that patrol axons; demyelination occurs in areas of intimate contact. PGL-1 confers this neurotoxic response on macrophages: macrophages infected with *M. marinum* expressing PGL-1 also damage axons. PGL-1 induces nitric oxide synthase in infected macrophages, and the resultant increase in reactive nitrogen species damages axons by injuring their mitochondria and inducing demyelination. Our findings implicate the response of innate macrophages to *M. leprae* PGL-1 in initiating nerve damage in leprosy.

## INTRODUCTION

Leprosy, like tuberculosis, presents as a granulomatous disease. These granulomas are usually cutaneous, reflecting the (∼30°C) growth optimum of *M. leprae*, similar to that of the human skin (∼34°C) (Bierman, 1936; Renault and Ernst, 2017; Truman and Krahenbuhl, 2001). *M. leprae* is the only mycobacterial infection that causes widespread demyelinating neuropathy, which results in the main morbidities of leprosy, including autoamputation of digits and blindness (Renault and Ernst, 2017; Scollard et al., 2006). Understanding the pathogenesis of leprosy neuropathy has been stymied by the inability to culture *M. leprae*, which has undergone severe reductive evolution of its genome to become an obligate intracellular pathogen (Cole et al., 2001; Renault and Ernst, 2017; Scollard et al., 2006). The lack of genetic tools for studying *M. leprae* is compounded by the lack of genetically tractable animal models that mimic the human disease. The athymic mouse footpad is used to grow *M. leprae* for research purposes, but the mouse does not manifest neurological disease (Scollard et al., 2006). While the nine-banded armadillo develops *M. leprae-*induced neuropathy, it suffers from a paucity of molecular and genetic tools (Scollard, 2008; Truman et al., 2014). Consequently, our understanding of the pathogenesis of leprosy neuropathy *in vivo* comes largely from studies of patients; however, the 4-10 year delay in the onset of symptoms precludes studies of the early events that lead to neuropathy (Noordeen, 1994).

Leprosy can present as a clinical spectrum; at the poles of this spectrum are paucibacillary (or tuberculoid) and multibacillary (or lepromatous) disease (Scollard 2006). The former is characterized by a vigorous immune response, while the latter, an ineffective one (Scollard 2006). Neuropathy features prominently in both forms of the disease. Hence, both bacterial determinants and host immune responses likely play roles in leprosy neuropathy, although the relative importance and mechanisms by which each contributes to nerve injury is poorly understood. *In vitro* studies suggest a model wherein *M. leprae* directly causes demyelination by infecting and dedifferentiating the Schwann cells that myelinate peripheral nerves (Rambukkana et al., 2002; Truman et al., 2014). These studies identified an *M. leprae* outer membrane lipid, phenolic glycolipid 1 (PGL-1), as critical for binding to laminin α2, an interaction thought to promote infection of the Schwann cells (Ng et al., 2000). However, this model fails to explain the neuropathy in paucibacillary leprosy, in which bacteria are seldom observed within nerve lesions (Shetty and Antia, 1996). Rather, a pathogenic CD4 T cell response, possibly acting through secreted cytokines, is implicated in the disease process (Renault and Ernst, 2017; Spierings et al., 2001). Further, the specific contributions of macrophages in leprosy neuropathy are unknown, although they are commonly infected and almost universally present in affected nerves (Job, 1973; Shetty and Antia, 1996).

The developing zebrafish is an effective model for studying mycobacterial pathogenesis using *M. marinum*, a close genetic relative of the *M. tuberculosis* complex and the agent of fish tuberculosis (Ramakrishan, 2004). The genetic tractability of the zebrafish, coupled with the optical transparency of its larva, allows host-bacterium interactions to be monitored in real-time, providing critical insights into disease pathogenesis (Cambier et al., 2014; Davis et al., 2002; Pagán and Ramakrishnan, 2015; Ramakrishan, 2004). Furthermore, adaptive immunity is not yet present at the larval developmental stage, permitting study of host-pathogen interactions in the sole context of innate immunity (Davis et al., 2002). Here, we exploit the optical transparency of larval zebrafish to directly visualize the earliest interactions of *M. leprae* with macrophages (Davis et al., 2002), and the initial events in nerve injury (Czopka, 2016; Villegas et al., 2012). We use *M. marinum* as a comparator for these studies because, like *M. leprae*, it grows at ∼30°C and produces cutaneous granulomatous infections in humans (Ramakrishan, 2004). However, it does not cause neuropathy. Our studies reveal that *M. leprae* interacts with macrophages and incites granulomas similar to *M. marinum*, but is unique in its ability to produce demyelination and axonal damage. We show that the innate macrophage response to PGL-1 triggers demyelination *in vivo*, even before bacilli have infected the glia. Finally, we determine the mechanism of nerve damage using *M. leprae* and *M. marinum* engineered to synthesize PGL-1.

## RESULTS

### *M. leprae* elicits typical responses in zebrafish larval macrophages

To determine if zebrafish larvae might be a useful model for studying early *M. leprae* infection, we first examined the earliest interactions of *M. leprae* with phagocytes, by injecting bacteria into the caudal vein or the hindbrain ventricle (Figure 1A), where phagocytes are rarely observed in the absence of infection (Davis et al., 2002). Aggregates of infected macrophages formed within four days (Figure 1B), similar to the case with *M. marinum* infection (Davis et al., 2002). Phagocyte recruitment to *M. marinum* infection is unique in two respects: 1) neutrophils are not recruited to the initial site of infection (Yang et al., 2012), and 2) macrophage recruitment is independent of TLR-signaling, but dependent on the monocyte chemokine CCL2 and its receptor CCR2 (Cambier et al., 2014). *M. leprae* shared both of these features with *M. marinum*: neutrophils were not recruited, while macrophages were (Figures 1C and 1D). Further, this recruitment was TLR/MyD88-independent and CCL2/CCR2-dependent (Figure 1D) (Cambier et al., 2014). The *M. marinum* phenolic glycolipid (PGL-mar) induces CCL2 expression and mediates CCL2/CCR2-dependent macrophage recruitment (Cambier et al., 2014), suggesting that PGL-1, may play a similar role in *M. leprae* infection.

**Figure 1.**
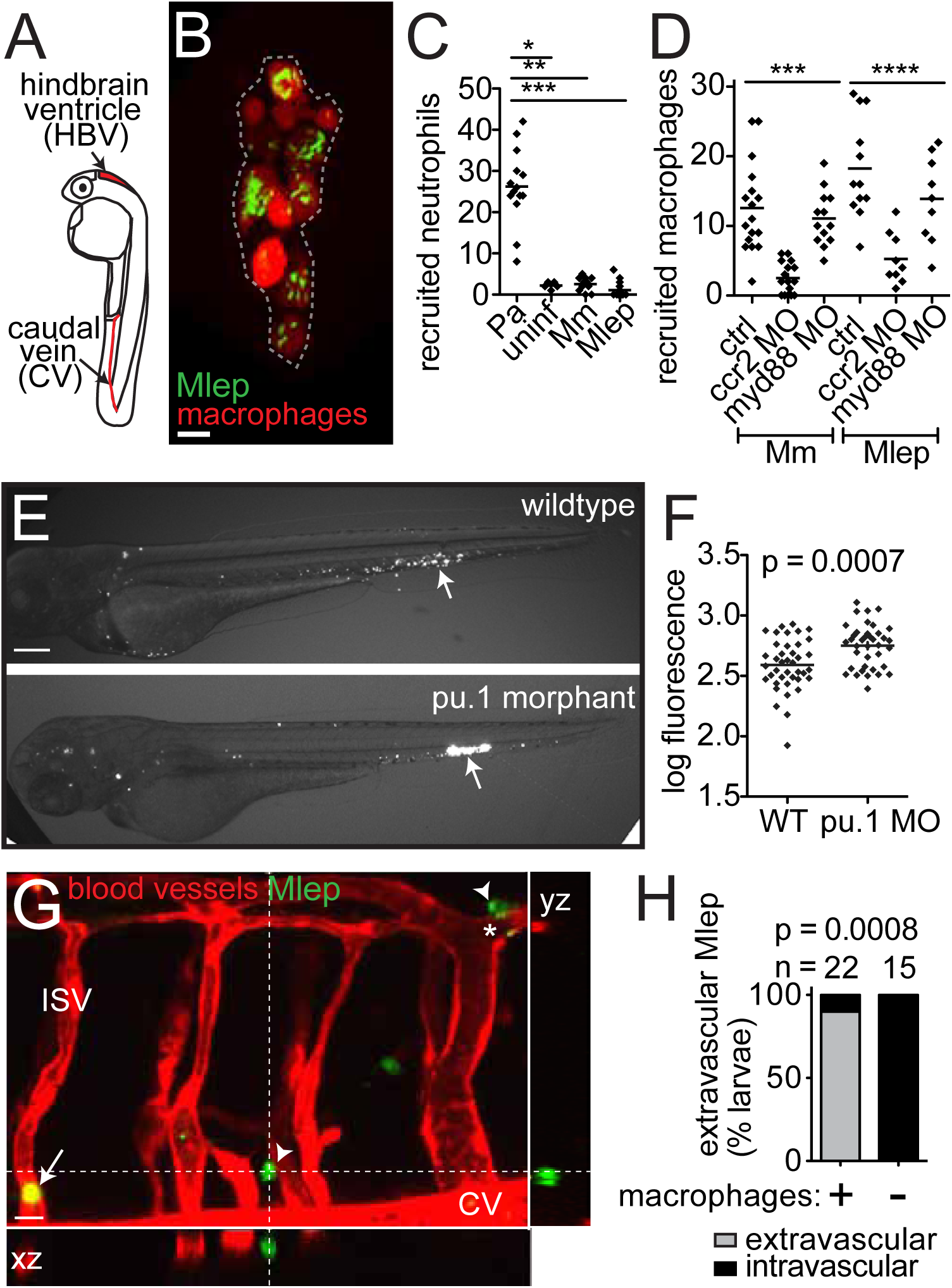
Early *M. leprae*-macrophage interactions are typical of other mycobacterial infections. (A) Diagram of larva 2 days post-fertilization (dpf) with injection sites indicated. (B) Representative confocal image of an early aggregate of fluorescent macrophages (dashed line), adjacent to the yolk sac, in an *mpeg1:YFP* larva at 4 days post-infection (dpi) with ∼10^4^ fluorescent *M. leprae* (Mlep). 10μm bar. (C) Mean number of neutrophils recruited to the hindbrain ventricle after injection of ∼100 colony-forming units (CFU) of *P. aeruginosa* (Pa), *M. marinum* (Mm), or Mlep in a 2 dpf larva; *p<0.05; **p<0.01; ***p<0.001 (one-way ANOVA with Bonferroni’s post-test). (D) Mean number of macrophages recruited to Mm or Mlep injection, as in C, in wildtype (WT) animals, or those made deficient in CCR2 or MyD88 by morpholino (MO) injection; ***p<0.001 (one-way ANOVA with Bonferroni’s post-test). (E) Representative fluorescent images of 2 dpi wildtype or macrophage-deficient PU.1 morphant larvae, infected with fluorescent Mlep as in B; the arrow indicates the injection site. 100μm bar. (F) Mean bacterial burden of larvae in E; unpaired Student’s t test. (G) Representative confocal image of the fluorescent vasculature of a 2 dpi *kdrl:dsRed* larva infected with fluorescent Mlep; bacteria reside within macrophages, apparent by Nomarski microscopy (Figure S1). The arrow indicates Mlep retained within vessels; arrowheads indicate Mlep outside of vessels; ISV, intersegmental vessel; asterisk, Mlep-infected macrophage surrounding the abluminal surface of the vessel. (H) Proportion of larvae in G with Mlep disseminated outside or contained within the vasculature, 4 days after caudal vein infection, in larvae depleted of macrophages, or not, by clondronate injection (Fisher’s exact test). N = number of larvae per group; all data representative of at least three separate experiments. See also Figure S1.

Macrophages play a dichotomous role in controlling *M. marinum* infection – they restrict bacterial numbers while promoting dissemination of bacteria from the infection site into deeper tissues (Clay et al., 2007). Similar to the case observed for *M. marinum, M. leprae*-infected animals depleted of macrophages (Clay et al., 2007) displayed higher bacterial burdens (Figure 1E and 1F). In addition, we assessed the role of macrophages in *M. leprae* dissemination, by infecting animals with fluorescent vascular endothelial cells (*kdrl:dsRed*). By 2 days post-infection (dpi), *M. leprae* escaped the vasculature and entered peripheral tissues in the majority of wildtype but not in macrophage-depleted larvae (Figure 1G and 1H). Furthermore, in wildtype animals, all *M. leprae* resided in macrophages (apparent by Nomarski imaging in Figure S1), suggesting these cells carried *M. leprae* from the circulation into tissues. In sum, *M. leprae* displays interactions with macrophages, from initial recruitment through granuloma formation, that resemble those seen for *M. marinum*. The presence of *M. leprae*-infected macrophages in the circulation of larvae mirrors findings in in human leprosy (Drutz et al., 1972).

### *M. leprae* infection alters myelin structure

We next investigated the interactions of *M. leprae* with cells of the zebrafish nervous system, to determine if infection produced demyelination. Transgenic *mbp* larvae express membrane-localized GFP that labels the myelinating membrane of glia in both the peripheral nervous system (Schwann cells) and central nervous system (oligodendrocytes) (Jung et al., 2010). Oligodendrocytes express all Schwann cell determinants that have been reported to interact with *M. leprae* (Table S1), and myelin structure is similar in the central and peripheral nervous systems (Morell and Quarles, 1999). Therefore, we studied *M. leprae* interactions with nerves in the spinal cord rather than peripheral nerves because of their easier accessibility. We injected fluorescent *M. leprae* into the dorsal spinal cord of larvae at 2-4 days post-fertilization (dpf), and imaged nerves at 4-8 dpf, a developmental stage at which these tracts have become myelinated (Czopka, 2016) (Figures 2A and 2B). At two dpi, we observed cellular protrusions from an otherwise intact myelin sheath, clustered around *M. leprae* in the nerve (Figure 2C). *M. marinum* injected into the dorsal spinal cord did not alter the myelinating membrane structure, even though the *M. marinum* burdens at the at the injection sites were higher than those in *M. leprae* infections (Figures 2C-2E). The *M. leprae*-induced myelin protrusions increased in size and number with time but always remained next to the bacteria (Figure 2F). Three-dimensional rendering showed that protrusions were doughnut-shaped, not spherical, suggesting that these structures were not cell bodies but rather protrusions of myelinating membrane (Figure 2G and Movie S1).

**Figure 2.**
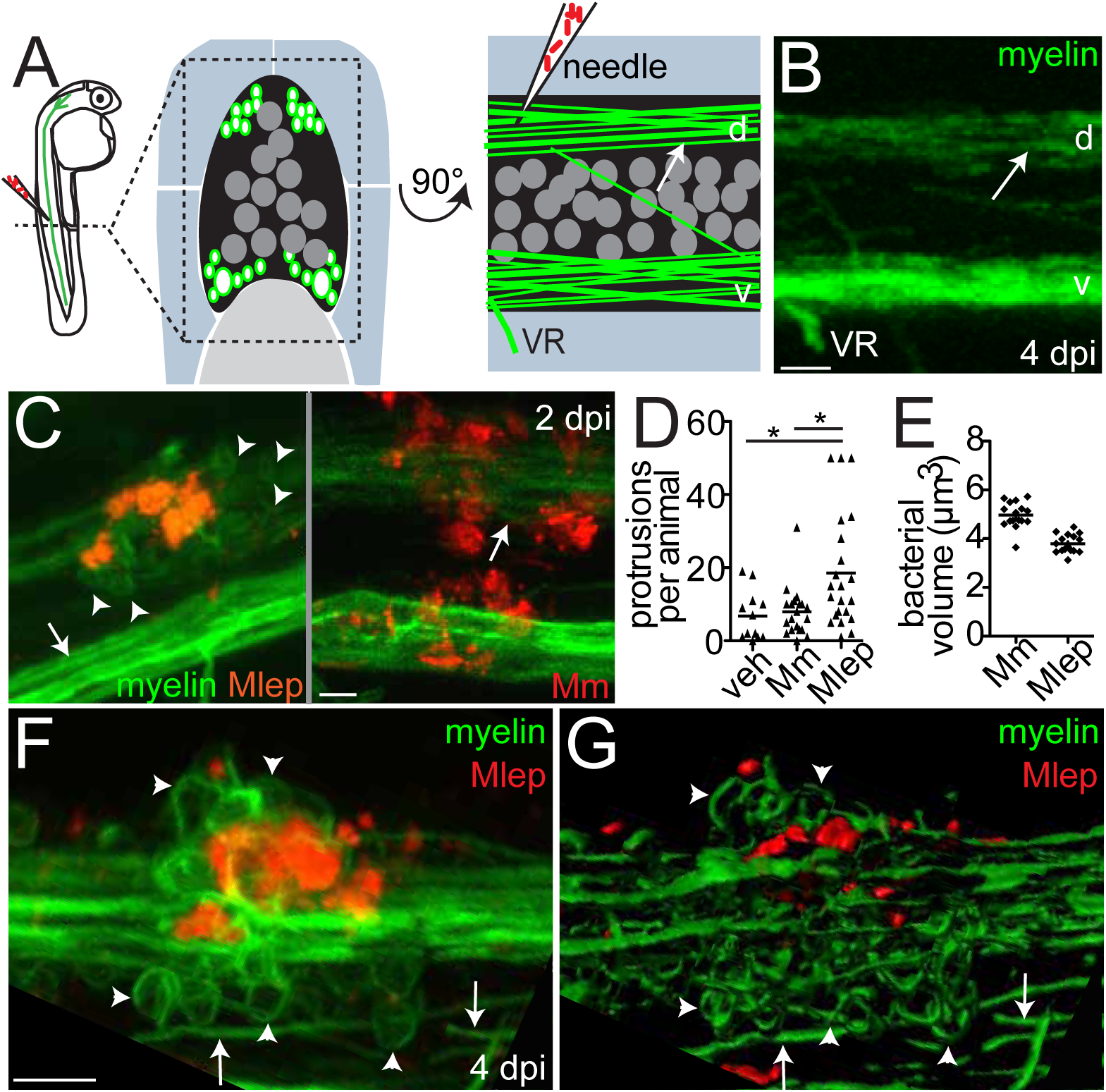
*M. leprae* triggers myelin dissociation. (A) Left, diagram of a spinal cord injection in an *mbp:eGFP-CAAX* larva (mbp), with fluorescent myelinating membrane, at 4 days post-fertilization (dpf). Transverse (middle) and sagittal (right) views of the region show the spinal cord (black), dorsal (d) and ventral (v) tracts of myelinated axons (green surrounding white axons), neuronal cell bodies (dark gray circles), and the ventral roots of spinal nerves (VR), surrounded by muscle (blue) and notochord (light gray). Arrows indicate intact myelin sheaths surrounding axons. (B) Confocal image corresponding to A. (C) Representative confocal images of *mbp* larvae, 2 days post-infection (dpi) with ∼10^4^ Mlep, or ∼200 colony-forming units (CFU) of *M. marinum* (Mm); arrowheads indicate myelin protrusions, quantified in D. (D) Mean number of myelin protrusions per animal upon injection with PBS vehicle (veh), Mm, or Mlep (one-way ANOVA, Bonferroni’s post-test). (E) Mean bacterial burden of larvae from C; measured by fluorescent pixel intensity as in Figure 1F. (F,G) Representative confocal image (F), and rendering (G), of myelin protrusions in a larva 4 dpi with ∼10^4^ Mlep (Movie S1).

### Expression of PGL-1 in *M. marinum* confers capacity to alter myelin structure

*In vitro* studies suggest that *M. leprae* interacts with glial determinants through a surface-localized long chain lipid, known as PGL-1 (*m/z* 2043.75), which carries a unique trisaccharide (Ng et al., 2000; Renault and Ernst, 2017) (Figure S2A). The phenolic glycolipid of *M. marinum* contains a monosaccharide and shorter lipid chains that renders it detectable at a lower mass value (*m/z* 1567.44) (Figures S2B and 3A). We wondered if the trisaccharide that is normally found on *M. leprae* PGL-1 would be sufficient to render *M. marinum* capable of altering myelin. We transformed *M. marinum* with the six *M. leprae* genes responsible for assembly of PGL-1’s terminal disaccharide (Tabouret et al., 2010). Ion chromatograms of total lipid from the transformant, *M. marinum:PGL-1*, proved that it produced triglycosylated PGL-1 (Figures S2C, 3A and 3B). PGL-1 expression conferred on *M. marinum* the ability to cause myelin protrusions, indistinguishable from those of *M. leprae* in both morphology and their invariable co-localization with the bacteria (Figures 3C-3E).

**Figure 3.**
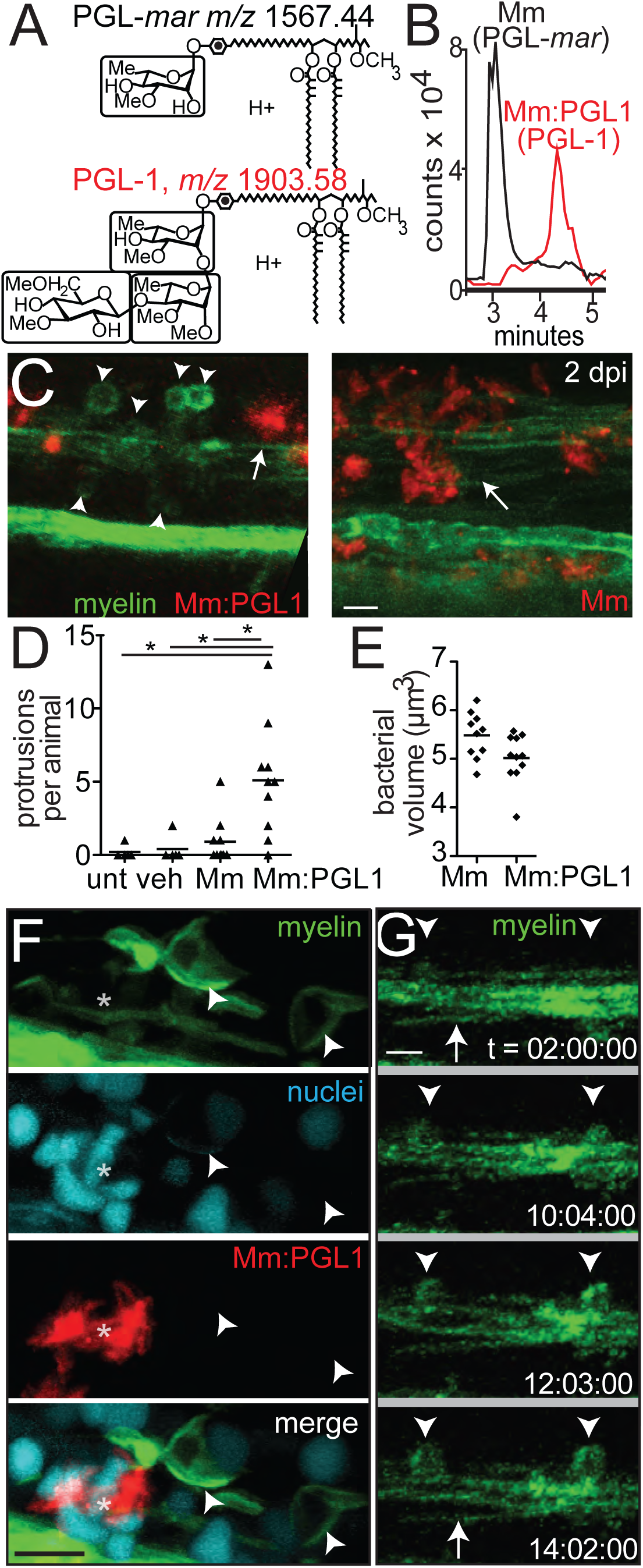
Phenolic glycolipid-1 triggers myelin dissociation. (A) Normal phase high performance liquid chromatography mass spectrometry measured at the known mass-to-charge ratios (*m/z)* for triglycosylated and monoglycosylated forms of PGL, leading to the separate detection of PGL-*mar* (*m/z* 1567.44, upper structure) and PGL-1 (*m/z* 1903.58, lower structure), as synthesized by recombinant *M. marinum:PGL-1* (Mm:PGL1). (B) Chromatograms of the ions depicted in A, showing the increased retention time of PGL-1 from Mm:PGL1 compared to that of PGL-*mar* from wildtype *M.marinum* (Mm). (C) Representative confocal images, as in Figure 2C, of 2 dpi larvae infected with ∼200 CFU Mm or Mm:PGL1; myelin protrusions quantified in D. (D) Mean number of myelin protrusions per animal in uninjected larvae (unt), or after injection with PBS vehicle (veh), Mm, or Mm:PGL1 (∼200 CFU each; one-way ANOVA with Bonferroni’s post-test). (E) Mean bacterial burden at the injection site of larvae from D. (F) Representative confocal image of a larva with fluorescently-labeled nuclei, 4 dpi with Mm:PGL1 (∼100 CFU). Asterisk indicates an aggregate of infected cells. (G) Stills from time lapse imaging of an *mbp* larva injected with Mm:PGL1, showing myelin protrusions forming from apparently intact myelin. Arrow, intact myelin sheath; arrowheads, myelin protrusions. Relative timecode; 10μm bars.

The protrusions, like those produced by *M. leprae*, did not colocalize with a histone marker that a labels cell nuclei. This suggested they did not simply represent an accumulation of oligodendrocyte cell bodies, but rather were composed of myelinating membrane (Figure 3F). Using time-lapse imaging to observe the formation of protrusions in real time, we observed that an intact myelin sheath near the *M. marinum:PGL-1* injection site began to condense and then bulge (Figure 3G). Protrusions formed by 10 hours post-infection, which expanded over time (Figure 3G). To further exclude the possibility that myelin protrusions represent recruitment or proliferation of oligodendrocyte cell bodies, we generated larvae with a single GFP-labeled oligodendrocyte. Time-lapse movies of these larvae showed that individual oligodendrocytes form myelin protrusions by retracting portions of myelinating membrane from previously intact sheaths (Movie S2A). This occurred after injection with *M. marinum:PGL-1*, but not with phosphate buffered saline (PBS) (compare Movies S2A and S2B). These findings strongly suggested that the protrusions arise from previously intact myelin sheaths, consistent with early demyelination. Similar to human leprosy (de Freitas and Said, 2013), myelin dissociation occurred in discrete areas, with the surrounding myelin sheaths remaining intact (Figure S3).

### Transmission electron microscopy shows PGL-1-mediated demyelination and axonal damage

Demyelination is best appreciated by transmission electron microscopy (TEM). We compared TEM images of transverse sections through areas of myelin protrusions at 2 days after infection to identical sections through the injection site of PBS-injected fish (Figures 4A-4C). TEMs from animals injected with *M. leprae* or *M. marinum:PGL-1* revealed a selective decrease in myelinated axons, while the total number of axons was preserved (Figures 4D and 4E; Figures S4A and S4B). Higher magnification images revealed apparently intact axons surrounded by disorganized myelin, with large spaces in between the individual lamellae (Figure S4C); this myelin decompaction is characteristic of early demyelination in human leprosy (Figure 4F) (Job, 1973; Shetty et al., 1988). The condensed, fragmented myelin, which was no longer associated with axons, was observed scattered throughout the extracellular space (Figures 4A-4C; Figure S4C).

**Figure 4.**
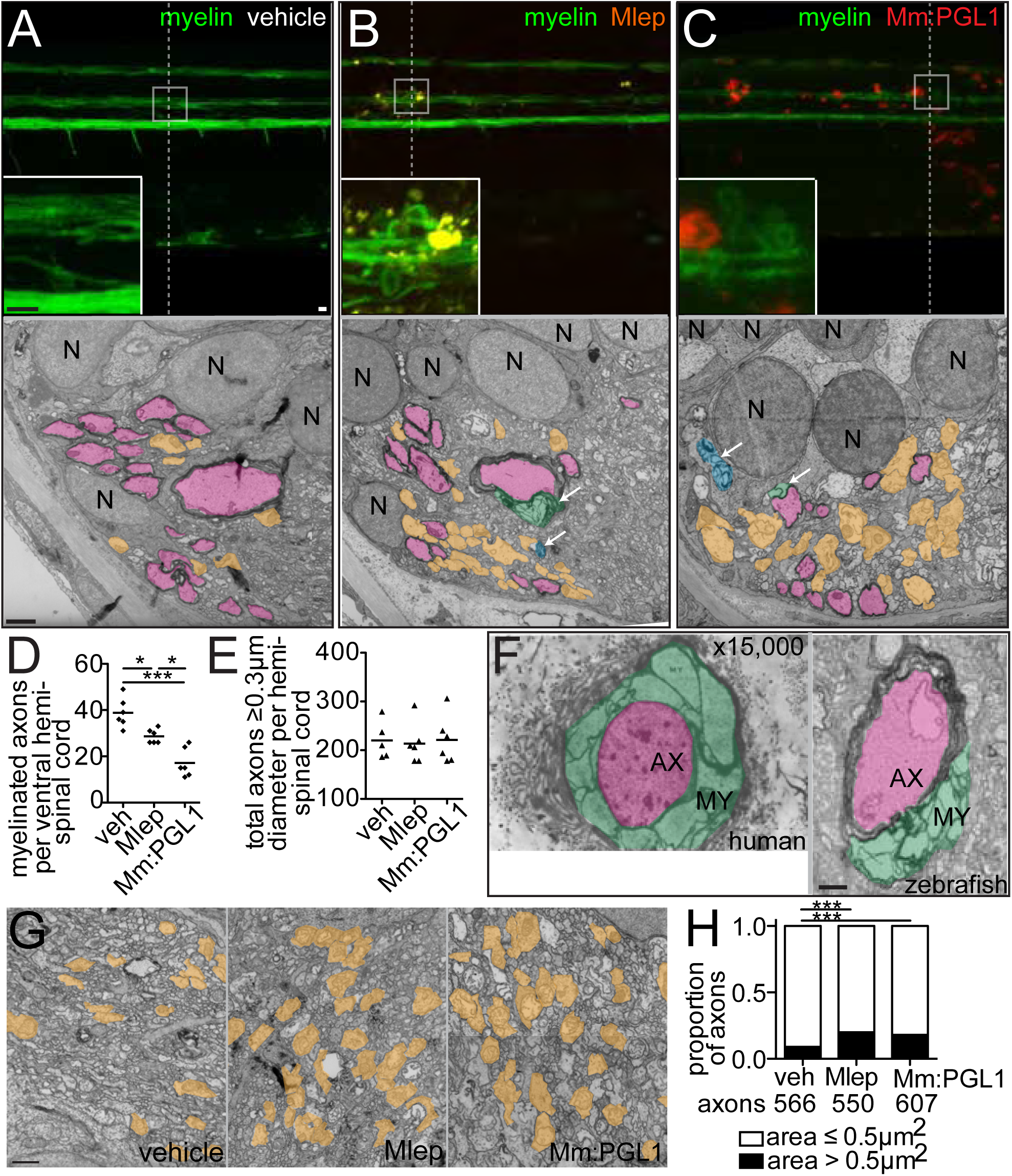
*M. leprae* alters nerve ultrastructure. (A-C) Representative confocal images of the spinal cord injection site (upper; 10μm bars) in *mbp* larvae at 2 dpi. Insets show magnifications of boxed regions; dashed lines indicate approximate location of the transmission electron micrograph (TEM) section, shown below with 1μm scale bars. Highlights indicate myelinated axons (pink), unmyelinated axons (orange), decompacted myelin (green highlight and arrows), and myelin dissociated from axons (blue highlight and arrows). N = neuronal cell body. (D,E) Mean number of myelinated axons (D) and total axons (E) per hemi-spinal cord section in larvae injected with PBS vehicle (veh), Mlep, or *M. marinum:PGL-1* (Mm:PGL1). (2 hemi-spinal cords scored per larvae; 3 larvae per group; one-way ANOVA with Bonferonni’s post test; *p<0.05; ***p<0.001). (F) Myelin decompaction in a radial nerve biopsy from a leprosy patient (left) (Job 1973, republished with permission), compared to similar alterations in the myelin of an Mlep-infected larva (right). MY = myelin; AX = axon; highlights indicate myelinated axons (pink) and decompacted myelin (green); 1μm scale bar. (G) TEMs of larvae obtained as in A, through matched anatomical regions. Nonmyelinated axons with diameter ≥ 0.5 μm^2^ are highlighted in orange; 1μm bar. (H) Proportion of nonmyelinated axons with area >0.5 or ≤ 0.5 μm^2^ from larvae obtained as in A. See also Figure S4.

*In vitro* studies have focused on *M. leprae*-induced demyelination as a mechanism of nerve injury (Rambukkana et al., 2002; Scollard, 2008). However, the peripheral neuropathy of human leprosy involves both myelinated and nonmyelinated axons (Medeiros et al., 2016; Shetty and Antia, 1996; Shetty et al., 1988). To test if nonmyelinated axons were also affected in zebrafish, we selected an area of the spinal cord containing only one myelinated axon surrounded by many nonmyelinated axons. We observed swelling of nonmyelinated axons, as evidenced by their increased area compared to PBS-injected control (Figures 4G and 4H). Thus, *M. leprae* and *M. marinum:PGL-1* rapidly induce damage to both myelinated and nonmyelinated axons in the zebrafish, similar to the pathological changes found in human leprosy.

### *M. leprae-induced* nerve damage is mediated by macrophages

Contrary to the previous model (Rambukkana, 2000), our findings *in vivo* did not support contact or infection of glia by *M. leprae* early in infection. We did not observe mycobacteria within myelin protrusions by confocal microscopy (Figure 2F), nor did we observe bacteria in direct contact with myelin or infected glia by TEM. All observed bacteria were within phagosomes of macrophages abutting the axons (Figures 5A and 5B). Given the presence of macrophages in the demyelinating lesions, we wondered if infected macrophages, rather than bacteria directly, initiated demyelination and nerve damage. Three findings in human leprosy support this idea: 1) macrophages, including those harboring *M. leprae*, are abundant in affected nerves even early in disease (Job, 1973; Nishiura, 1960; Pandya and Antia, 1974; Shetty and Antia, 1996; Shetty et al., 1988). 2) Early stages of demyelination feature vacuolar myelin structures, in which the lamellae have split and separated (Job, 1973), associated with infected macrophages beneath the basement membrane of Schwann cells. 3) The unique trisaccharide of *M. leprae* PGL-1 confers both demyelinating (Ng et al., 2000; Renault and Ernst, 2017) and macrophage-modulating effects *in vitro* (Fallows et al., 2016; Manca et al., 2011; Tabouret et al., 2010). The plausibility of a macrophage-induced mechanism is further supported by findings that macrophages mediate demyelination and nerve damage in multiple sclerosis and Guillain-Barré syndrome (Bogie et al., 2014; Martini et al., 2008; Martini and Willison, 2016; Nikic et al., 2011).

**Figure 5.**
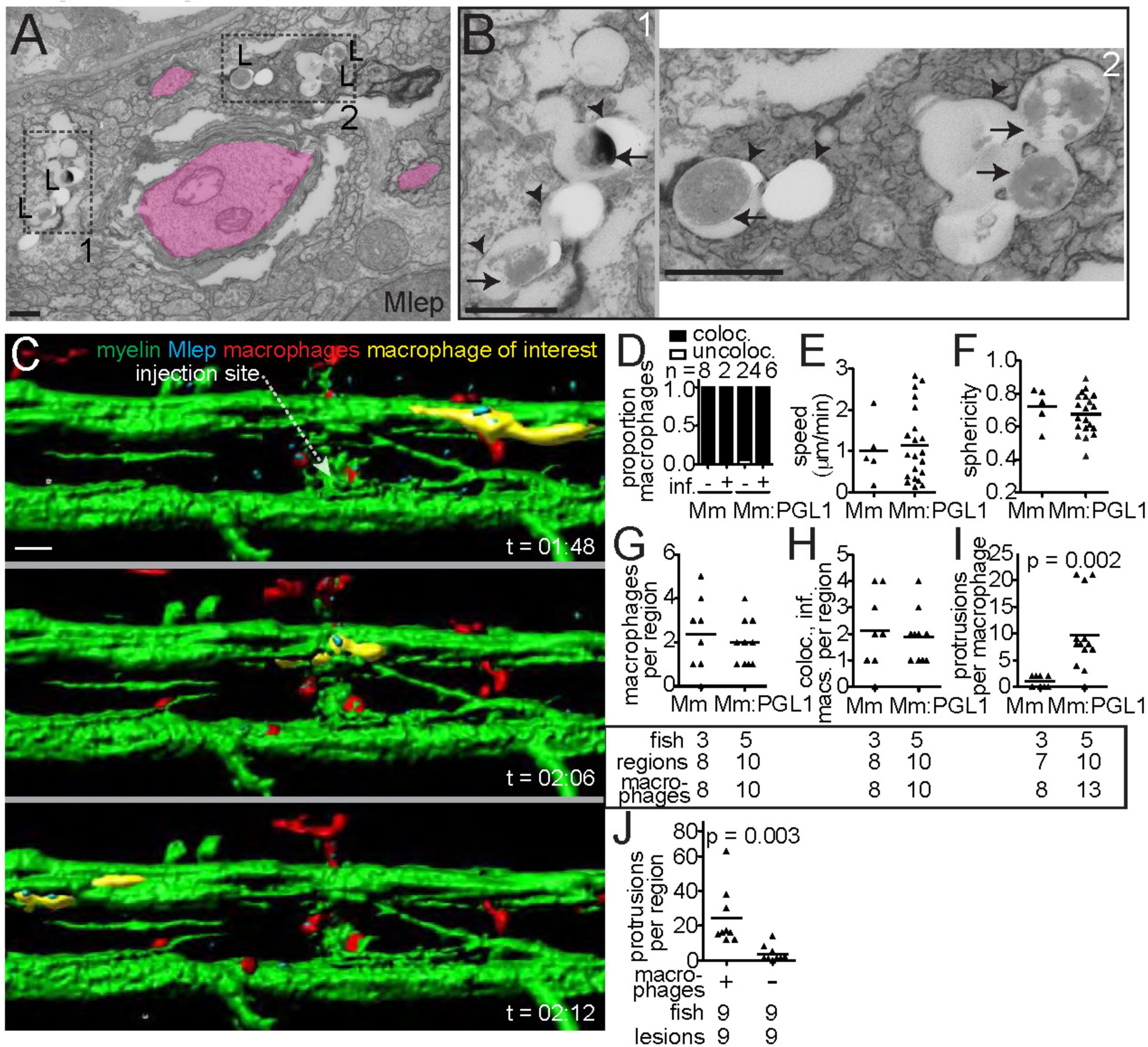
Macrophages mediate *M. leprae* demyelination. (A) TEM from 6 dpi larva showing Mlep bacilli (L) within a phagocyte contacting a myelinated axon. Dashed line indicated insets 1 and 2, shown in B; myelinated axons highlighted in pink; 1μm bar. (B) Insets from A, showing the Mlep double membrane (arrows) and phagosomal membranes (arrowheads); 1μm bar. (C) Rendered still images from a timelapse movie (Movie S4) of an Mlep-infected double transgenic *mbp;mpeg1* larva, with fluorescent macrophages and myelinating membrane. At 4 days post-fertilization, the larva was infected in the spinal cord and immediately imaged for 12 hours, revealing infected macrophages patrolling intact myelin sheaths (a myelin-patrolling, infected macrophage highlighted in yellow). 10μm bar. (D) Proportion of uninfected (−) or infected (+) macrophages that colocalized with myelin, in 4 dpf larvae infected with *M. marinum* or *M. marinum:PGL-1*. n = number of macrophages scored. (E) Mean speed of macrophages in the larvae from D. (F) Mean sphericity of macrophages in the larvae from D. (G) Mean number of macrophages per infected region, in *mbp* larvae 2 dpi with *M. marinum* or *M. marinum:PGL-1*. Numbers of fish, lesions, and macrophages scored per group are indicated. (H) Mean number of macrophages per region that were both infected and myelin-colocalized, in the larvae from G. (I) Mean number of myelin protrusions per macrophage, in the larvae from G. Student’s t-test. (J) Myelin protrusions per Mlep-infected region, in wildtype *mbp* larvae (+) or those depleted of macrophages (−) by injection with *irf8* morpholino and lipo-clodronate. Student’s t-test. Data representative of at least two separate experiments. See also Movies S4-S6.

Macrophages are associated with nerves under homeostatic conditions in humans and rodents, both in the peripheral and central nervous systems (Kierdorf et al., 2013; Klein and Martini, 2016; Müller et al., 2010). In the case of nerve injury, their numbers increase (Klein and Martini, 2016), presumably because they play roles in scavenging debris and repair. In the zebrafish too, we observed macrophages arriving from the blood and patrolling axons in uninjected larvae, and their numbers increased in response to the trauma of PBS injection (Movies S3-S5).

We asked if infection with PGL-1-expressing bacteria made these macrophages capable of demyelinating axons. We used blue or far-red fluorescent bacteria to infect transgenic larvae with green fluorescent myelinating membrane and red fluorescent macrophages. Immediately after infection, macrophages were recruited to the injection site, entered the spinal cord, and phagocytosed the majority of the bacteria; this was equally the case for *M. leprae, M. marinum* and *M. marinum:PGL-1* (Movies S4-S6). Moreover, in the context of each infection, macrophages, whether infected or not, patrolled the axons, assuming a flattened, elongated shape as they moved between them (Figure 5C and Movies S6-S8). We noted that some infected macrophages moved more slowly and eventually became sessile within the first twelve hours, resulting in prolonged intimate contact with the myelin in discrete areas. This slowing down of infected macrophages has been noted in *M. marinum* granulomas (Davis and Ramakrishnan, 2009). Here too, we observed more slowly moving infected macrophages in the context of all three infections, suggesting it was an infection- but not PGL-1-dependent phenomenon (Movies S6-S8). This was confirmed by a quantitative comparison of macrophage behavior during the first twelve hours following *M. marinum* versus *M. marinum:PGL-1* infection: there were no differences in their speed, shape or tendency to associate with myelin (Figures 5D-5F). Macrophage co-localization with myelin continued to be similar between the two bacterial groups at two days post-infection, when demyelination begins (Figures 5G and 5H). Yet, only colocalization of *M. marinum:PGL-1-infected* macrophages with myelin produced myelin protrusions (Figure 5I). All demyelinating lesions were associated with macrophages in 10 of 11 animals scored (Table S2; p = 0.01, two-tailed binomial test with an expected 0.5 frequency). In the eleventh animal, two of the three demyelinating lesions were associated with macrophages, while the third had defined clusters of bacteria with residual fluorescent macrophage membrane, suggesting that the co-localized, infected macrophage had died (Figure S5). Finally, to directly test if macrophages were required for *M. leprae*-induced demyelination, we created macrophage-depleted fish by administering an *irf8* morpholino followed by clodronate liposomes (Pagán et al., 2015). Macrophage depletion reduced myelin protrusions by 85% in *M. leprae*-infected larvae, confirming the essential role of macrophages in early demyelination (Figure 5J).

### A theoretical framework for the mechanism of PGL-1- and macrophage-dependent demyelination

The peripheral neuropathy of leprosy is analogous to that of Gullain Barré syndrome, in which the production of macrophage effectors, including reactive oxygen and nitrogen species (ROS and RNS), is implicated in nerve damage (Kiefer et al., 2001; Martini and Willison, 2016; Nikic et al., 2011). However, the mechanisms by which macrophages produce these effectors and mediate nerve damage have been more extensively described for multiple sclerosis, a demyelinating disease of the central nervous system (Bogie et al., 2014; Nikic et al., 2011). As with leprosy, multiple sclerosis affects both myelinated and nonmyelinated axons. In multiple sclerosis, it has been suggested that macrophage-derived ROS/RNS can trigger mitochondrial pathology, initiating swelling and destruction of mitochondria and axons (Nikic et al., 2011). Similarly, the macrophages present in leprosy skin and nerve biopsies express inducible nitric oxide synthase (iNOS) and contain nitrotyrosine, the stable end product of nitric oxide and superoxide reaction with tyrosine residues (Lockwood et al., 2011; Schön et al., 2004; Teles et al., 2013). Moreover, recent work has shown that mitochondria are swollen and damaged in both myelinated and nonmyelinated axons (Medeiros et al., 2016). Together, these findings suggest a model in which PGL-1 induces iNOS expression in infected macrophages, resulting in oxidative damage to mitochondria of adjacent axons. This model generates three testable predictions: 1) PGL-1-expressing bacteria induce production of iNOS and nitric oxide in the macrophages they infect; 2) PGL-1-induced nerve damage is nitric oxide-dependent; and 3) nerve damage is linked to mitochondrial damage, which is also PGL-1-dependent.

### Nitric Oxide Production by Macrophages in Response to PGL-1 Mediates Demyelination

To test the first prediction of our model, we asked if PGL-1 induces *Nos2* (the gene that encodes iNOS) in cultured murine bone marrow-derived macrophages. *M. marinum:PGL-1* induced 2.8-fold more *Nos2* in macrophages than wildtype *M. marinum*, showing a substantial contribution from PGL-1 (Figure 6A). In the zebrafish too, *M. marinum:PGL-1*-infected macrophages were iNOS-and/or nitrotyrosine-positive, both in the periphery and the nervous system (Figures 6B, S6A and S6B). Again, *M. marinum:PGL-1* infection was associated with more iNOS- and nitrotyrosine-positive macrophages than wildtype *M. marinum* (Figures 6C and 6D). Thus, PGL-1-expressing mycobacteria induce macrophages to produce nitric oxide through transcriptional induction of iNOS.

**Figure 6.**
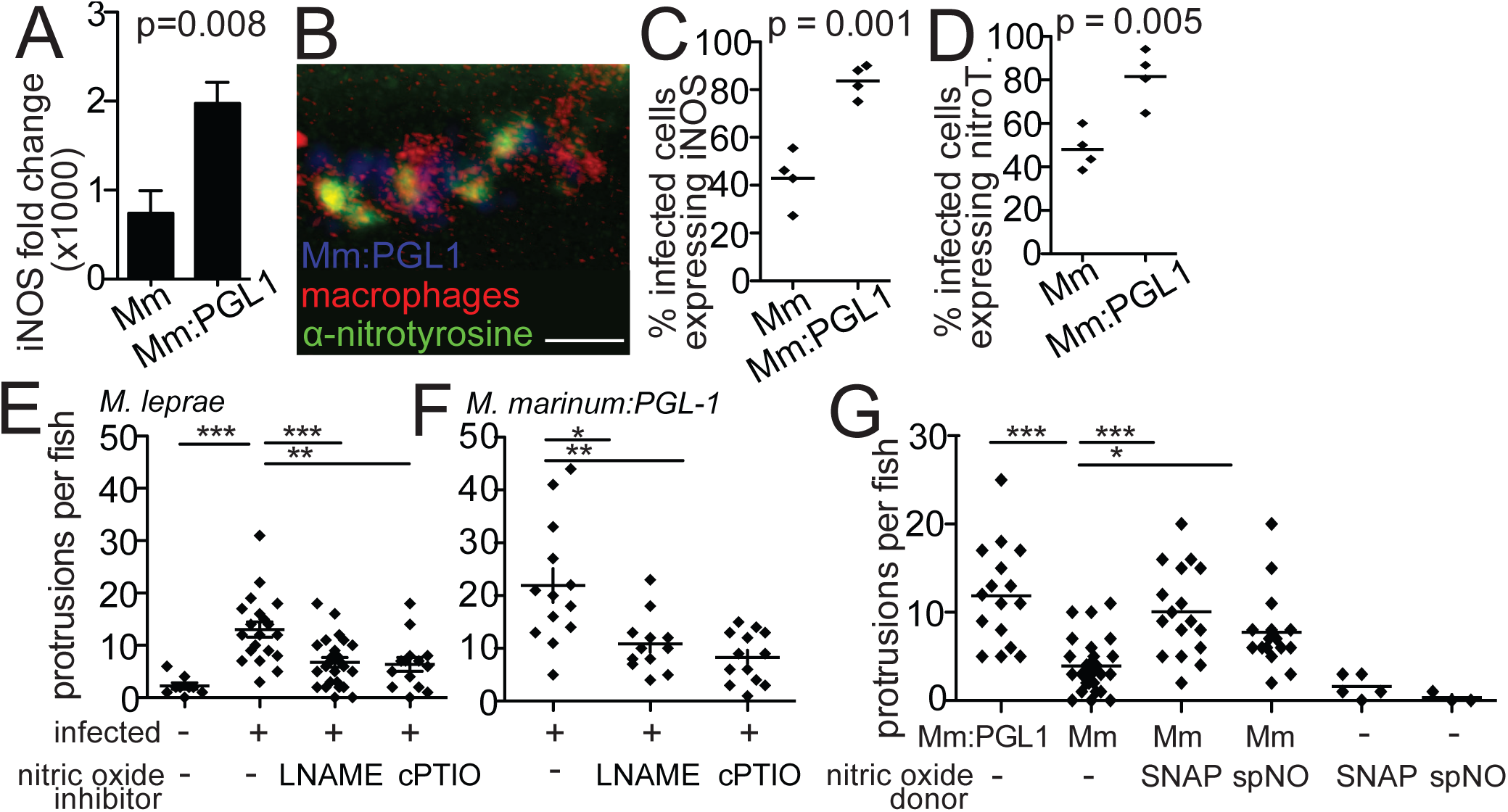
Nitric Oxide is Necessary for Early Demyelination. (A) Mean (±SEM) fold change of *Nos2* transcript in wildtype murine macrophages, 6 hours after infection with wildtype *M. marinum* or *M. marinum:PGL-1* (both MOI 1), compared to uninfected cells. (Average of 3 independent experiments; student’s t-test) (B) Representative confocal images of a macrophage aggregate in the spinal cord of an *mpeg1* larva infected with *M. marinum:PGL-1* and stained with α-nitrotyrosine antibody. 10μm bar. (C) Mean percent of infected, *mpeg1*-positive macrophages that also express iNOS, in larvae 5 dpi with wildtype *M. marinum* or *M. marinum:PGL-1*. (Student’s t-test) (D) Mean percent of infected, *mpeg1*-positive macrophages that stain with α-nitrotyrosine antibody (nitroT.), in larvae as in C. (Student’s t-test) (E) Mean number of myelin protrusions per animal in *mbp* larvae 2 dpi with *M. leprae*, which were treated with 0.5% DMSO vehicle (veh), iNOS inhibitor (L-NAME) or ROS/RNS scavenger (cPTIO). (**p<0.01; ***p<0.001; one-way ANOVA with Dunnett’s multiple comparison test) (F) Mean number of myelin protrusions per animal in *mbp* larvae 2 dpi with *M. marinum:PGL-1*, treated as in E. (G) Mean number of myelin protrusions per animal in larvae infected as in F, which were soaked post-injection in 0.5% DMSO vehicle (“-“) or in nitric oxide donors SNAP or spermine NONOate (spNO). (*p<0.05; ***p<0.001; one-way ANOVA with Dunnett’s multiple comparison test). See also Figure S6.

To test the second prediction of our model, we asked if nitric oxide induces early demyelination by treating infected fish with the iNOS inhibitor L-NAME or the ROS/RNS scavenger cPTIO. Both treatments inhibited demyelination, in larvae infected with *M. leprae* or *M. marinum:PGL-1* (Figures 6E and 6F). Conversely, the nitric oxide donors SNAP and spermine NONOate induced demyelination in larvae infected with wildtype *M. marinum*, but nitric oxide donors failed to cause demyelination if *M. marinum* was not present (Figure 6G), suggesting that nitric oxide acts with other determinants induced by virulent mycobacteria, independent of PGL-1 (Figure 6G).

### PGL1-induced axonal damage is associated with mitochondrial swelling and loss

The third prediction of our model is that mitochondrial damage is linked to nerve damage, and is dependent on PGL-1 production by bacteria. Confocal microscopy of *mbp* larvae expressing a fluorescent protein in axonal mitochondria revealed both mitochondrial swelling and selective loss in regions close to demyelinating lesions (Figure 7A). TEMs through demyelinating lesions of *M. leprae* and *M. marinum:PGL-1*-infected larvae had fewer axonal mitochondria compared to PBS-injected larvae (Figures 7B and 7C). The remaining mitochondria were enlarged in infected larvae compared to PBS-injected controls, similar to the mitochondrial swelling reported for leprosy and multiple sclerosis (Medeiros et al., 2016; Nikic et al., 2011) (Figures 7D and 7E). If mitochondrial damage is linked to axonal damage, then it should be most prevalent in swollen axons. Two analyses showed that this was the case: first, the increase in mitochondrial area in infection over PBS-control occurred in axons with an area ≥ 0.5 μm^2^, but not in those with an area < 0.5 μm^2^ (Figures 7F and 7G). Second, within each of the three cohorts, mitochondrial area was increased only in the large axons (≥ 0.5 μm^2^) of *M. leprae* and *M. marinum:PGL-1*-infected larvae, not in PBS-injected larvae (Figure 7H). As expected, there was no difference in mitochondrial area in the axons of PBS-treated animals, where the differences in axon size reflect normal physiological variation, rather than pathology. Collectively, these findings support the model that that reactive nitrogen species produced by infected macrophages damage axonal mitochondria and cause demyelination.

**Figure 7.**
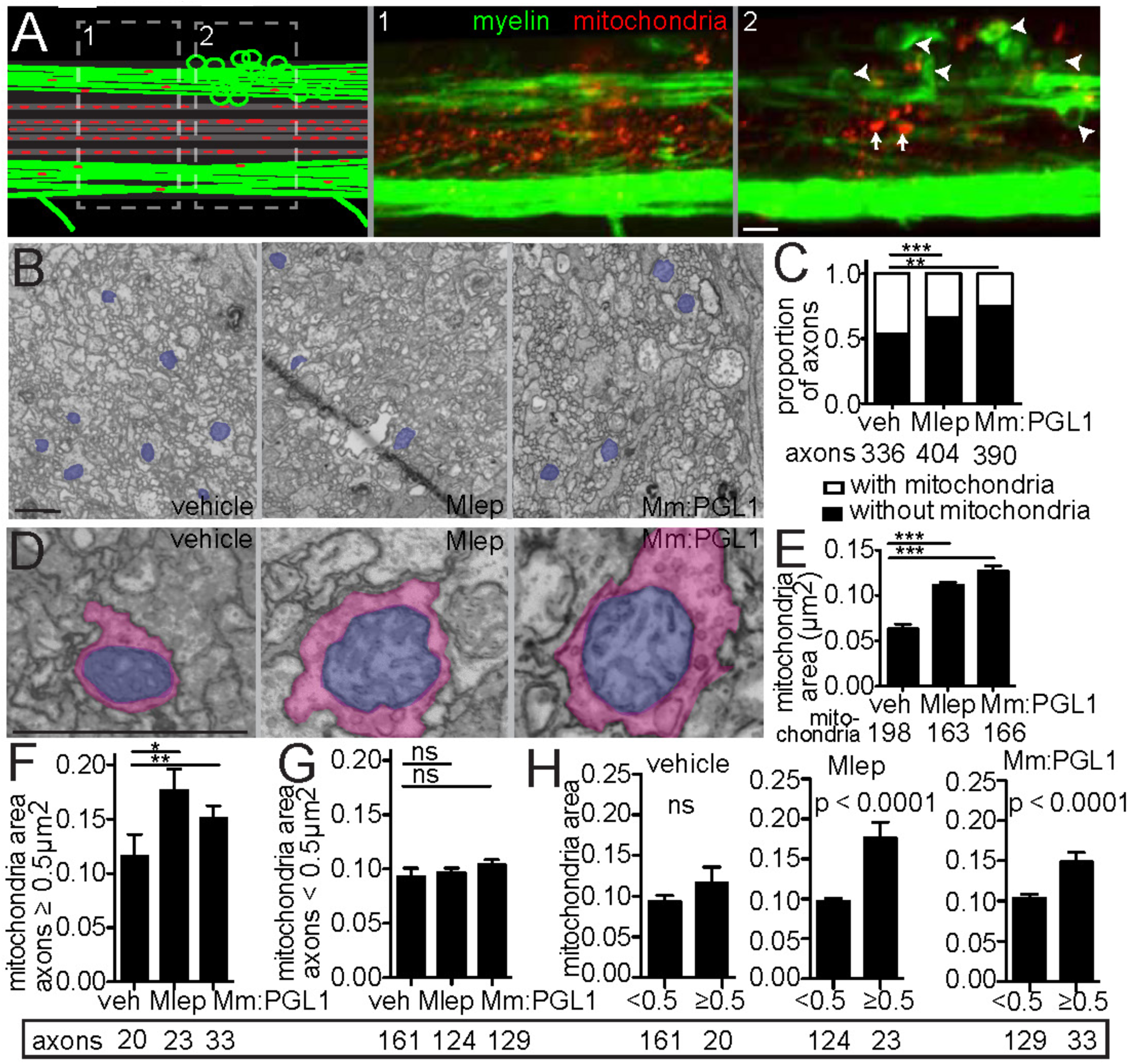
*M. leprae* infection damages axon mitochondria. (A) Diagram (left) of a demyelinating lesion in a 2 dpi *mbp* larva with fluorescent mitochondria in axons. Dashed boxes indicate insets 1 and 2, with corresponding confocal images, showing mitochondria outside the lesion (inset 1) and those within the lesion (inset 2). Arrowheads indicate myelin protrusions; arrows indicate enlarged mitochondria. (B) Representative TEMs of matched anatomical regions showing the number of axon mitochondria (blue) in larvae injected with PBS, Mlep or *M. marinum:PGL-1*. 1μm bar. (C) Proportion of axons with mitochondria to those without, in larvae as in B; numbers of axons scored listed below (contingency analysis corrected for multiple comparisons; **p = 0.004; ***p<0.0002). (D) Representative TEMs of enlarged mitochondria (blue) within enlarged axons (pink), in larvae as in B. 1μm bar. (E) Mean (±SEM) area of mitochondria in axons, in larvae as in B; number of mitochondria scored are listed below. (***p<0.001; one-way ANOVA with Dunnett’s multiple comparison). (F) Mean (±SEM) area of mitochondria in nonmyelinated axons with area ≥ 0.5 m^2^, in larvae as in B (*p<0.05, **p<0.01; one-way ANOVA with Dunnett’s multiple comparison). (G) Mean (±SEM) area of mitochondria in nonmyelinated axons with area < 0.5 m^2^, in larvae as in B. (H) Data from F and G displayed per experimental group, showing mean (±SEM) area of mitochondria in large versus small nonmyelinated axons (one-way ANOVA with Dunnett’s multiple comparison).

## DISCUSSION

Our work suggests a mechanism for the earliest nerve injury associated with leprosy: over-exuberant production of nitric oxide by macrophages, in response to the *M. leprae*-specific PGL-1, damages axonal mitochondria and initiates demyelination (Figure 8A). Mycobacterial phenolic glycolipids likely evolved to increase infectivity by recruiting macrophage subsets that are particularly permissive to mycobacterial infection (Cambier et al., 2014). We find here that the specialized, triglycosylated form of *M. leprae* PGL-1 retains this basal role, while acquiring additional macrophage-modulating functions. PGL-1 has been found to alter inflammatory mediator expression in cultured macrophages (Fallows et al., 2016; Manca et al., 2011; Tabouret et al., 2010), and our work now assigns a central role for this immunomodulation in early leprosy neuropathy.

**Figure 8.**
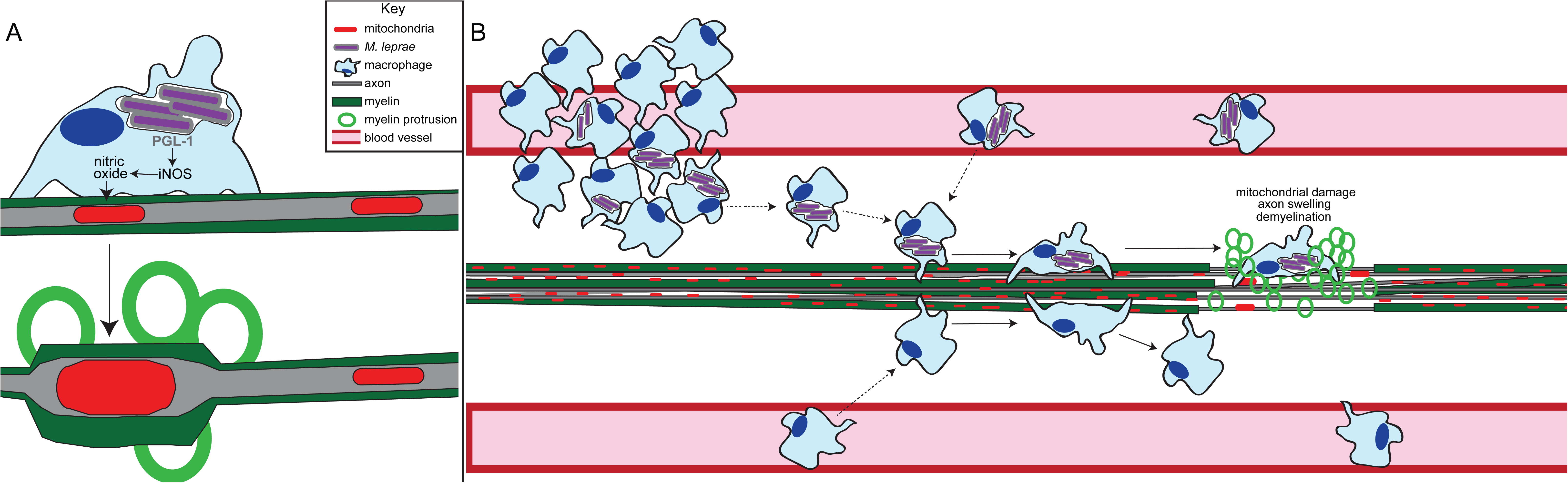
Macrophage-mediated demyelination in leprosy. (A) Macrophages upregulate production of iNOS in response to the *M. leprae* surface lipid, PGL-1. iNOS produces nitric oxide, which damages mitochondria and causes demyelination in nearby axons. (B) Circulating *M. leprae*-infected macrophages reach axons by exiting the perineural blood vessels, which supply peripheral nerves, or by arriving cutaneously from a skin lesion that overlies a nerve. Once there, infected macrophages patrol axons and occasionally become sessile. In response to the PGL-1 synthesized by their infecting bacilli, these sedentary macrophages cause nerve injury characterized by axon and mitochondrial swelling, demyelination, and mitochondrial loss. If the macrophage dies, it releases *M. leprae* into the extracellular environment of the nerve, where they can directly infect Schwann cells.

In terms of the relevance of our findings to human leprosy, macrophages, often infected, are a consistent presence within early (often clinically asymptomatic) nerve lesions of leprosy patients (Job, 1973; Nishiura, 1960; Pandya and Antia, 1974; Shetty and Antia, 1996; Shetty et al., 1988). As to how infected macrophages might reach nerves, one way is by direct seeding from a skin granuloma into an underlying nerve trunk. In support of this possibility, a leprosy cohort study found that the most significant risk factor for development of neuropathy in a peripheral nerve was the presence of an overlying skin lesion (Van Brakel et al., 2005). A second possibility is through hematogenous dissemination. This work and others suggest that circulating macrophages patrol axons in both the peripheral and central nervous systems under homeostatic conditions, reaching axons by extravasating from local blood vessels (Kierdorf et al., 2013; Klein and Martini, 2016; Müller et al., 2010)(Figure 8B). Bacteremia is common in leprosy (Drutz et al., 1972; Ganapati and Chulawala, 1976; J. E. Lane et al., 2006; Kaur et al., 1993). Bacteria have been observed in the blood vessels of apparently normal skin of leprosy patients (Ganapati and Chulawala, 1976), and many of the bacteria in the blood are in mononuclear phagocytes (Drutz et al., 1972). We suggest that infected macrophages have a similar propensity to reach nerves and patrol them to their uninfected counterparts. Some may slow down and stall as *Mycobacterium*-infected macrophages are wont to do (Davis and Ramakrishnan, 2009). The resultant prolonged intimate contact with the nerve may initiate damage through the mechanism we have uncovered (Figure 8B). This hematogenous dissemination model predicts that *M. leprae*-infected macrophages are widely distributed in nerves. Indeed, biopsies of the apparently normal skin of leprosy patients find subclinical, diffuse neuropathy in conjunction with infected macrophages (Ganapati et al., 1972; Pandya and Antia, 1974; Rea et al., 1975). Finally, household contacts of leprosy patients are significantly more likely to have *M. leprae* DNA in their peripheral blood than noncontacts, and longitudinal follow-up shows that these individuals are more likely to develop leprosy (Reis et al., 2014; Wen et al., 2013), suggesting that hematogenous dissemination of *M. leprae* is a very early and significant step in the pathogenesis of peripheral nerve damage.

Our finding that both myelinated and nonmyelinated axons are damaged by this mechanism further suggests its relevance to human leprosy neuropathy, which affects both types of axons (Medeiros et al., 2016; Shetty and Antia, 1996; Shetty et al., 1988). Moreover, nonmyelinated cutaneous nerve endings are often affected early in infection, even before neurological symptoms appear (de Freitas and Said, 2013; Ganapati et al., 1972; Pandya and Antia, 1974). The idea that demyelination is a pathological manifestation, rather than a cause of nerve injury, has gained traction in the context of other demyelinating diseases such as multiple sclerosis (Aboul-Enein et al., 2006; Nikic et al., 2011).

The prior model, which explains the neurotropism of *M. leprae* by evoking direct binding of PGL-1 to Schwann cell laminin α2 (Ng et al., 2000) is problematic in at least four ways. First, other mycobacterial species that lack PGL-1 and fail to cause neuropathy are, nevertheless, able to bind laminin α2 (Marques et al., 2001; Scollard, 2008). Second, the model does not explain how *M. leprae*, a nonmotile bacterium, reaches Schwann cells. Third, by requiring high bacterial burdens within Schwann cells to cause demyelination, the model fails to explain the clinical findings of nerve damage very early in infection, when only a few bacteria are present in nerve lesions. Fourth, the earliest nerve impairment in leprosy is in thermal sensation, which is mediated by nonmyelinated fibers (de Freitas and Said, 2013; Shetty and Antia, 1996; Van Brakel et al., 2003). Our findings and model resolve these inconsistencies, by showing that it is the PGL-1-stimulated innate macrophages that initiate damage to nerves regardless of their myelination. This early innate-immune mediated nerve injury may then progress by distinct mechanisms in multibacillary and paucibacillary leprosy (Cammer et al., 1978; Spierings et al., 2001; Wisniewski and Bloom, 1975). In the face of an inadequate adaptive immune response in multibacillary leprosy, the inability of macrophages to control bacterial growth may result in their death, releasing bacteria into the extracellular milieu of the nerve. These released bacteria could then be taken up by Schwann cells. In paucibacillary leprosy, the onset of an adaptive immune response may enable infected macrophages to control intracellular *M. leprae*, while further enabling, or even enhancing, their neuropathological response (Renault and Ernst, 2017; Scollard, 2008). This may be through the induction of pro-inflammatory cytokines such as interferon-γ (Teles et al., 2013), which may act by further stimulating reactive oxygen and nitrogen species, or by engaging distinct mechanisms.

Production of nitric oxide by macrophages and other myeloid cells has been implicated in mitochondrial dysfunction and subsequent axonal injury in multiple sclerosis and Guillain-Barré syndrome (Geleijns et al., 2007; Nikic et al., 2011). Our work may offer insights into these and other neurodegenerative diseases in which myeloid cells are increasingly recognized as contributing to neuropathology (Asiimwe et al., 2016; Thompson and Tsirka, 2017), as well as provide a new experimental system in which to explore them.

## ACKNOWLEDGEMENTS

Live *M. leprae* was regularly supplied by Ramanuj Lahiri and James Krahenbuhl at the National Hansen’s Disease Programs through the generous support of the American Leprosy Missions and Society of St. Lazarus of Jerusalem. We thank Stanley Falkow and Paul Edelstein for reading and editing the manuscript, David Raible, William Talbot and Paul Edelstein for discussion and advice, Paul Edelstein for help with statistics, Antonio Pagán for advice and help with the macrophage depletion experiments, Yuan Dong and James Cameron for zebrafish husbandry, William Talbot and Christophe Guilhot for reagents, Frank Ciampi for assistance with movies, and Marianne Cilluffo at the BRI Electron Microscopy Core for assistance with electron microscopy studies. Confocal imaging was performed at the CNSI Advanced Light Microscopy/Spectroscopy Facility at UCLA. This work was supported by an A.P. Giannini Foundation Postdoctoral Fellowship, NIH T32 AI1007411 and an NIH NRSA postdoctoral fellowship AI104240 (C.A.M.), an NSF Predoctoral Fellowship and NIH Bacterial Pathogenesis Training Grant Award (C.J.C.), a UCLA Clinical Translational Science Institute grant UL1TR001881 (K.K.S.), K08AR066545 (P.O.S.), U19 AI 111224 and R01 AI 049313 (D.B.M.), NIH R01AR064582 (A.S.), NIH R37AI054503, the NIH Director’s Pioneer Award, and a Wellcome Trust Principal Research Fellowship (L.R.).

## SUPPLEMENTAL FIGURE LEGENDS

**Supplemental Figure S1. *M. leprae*-infected macrophages escape the circulation.** Related to Figures 1G and 1H.

(A) Confocal image from Figure 1G of a *kdrl:dsRed* larva, which has fluorescent blood vessels, 2 days post-infection (dpi) with fluorescent *M. leprae* (Mlep); dashed lines define insets (1-4) shown in B.

(B) Monochannel and merged images of insets from A, showing Mlep within cells, likely macrophages, which are visible by Nomarski microscopy. Arrows, intracellular Mlep retained in vessels; arrowheads, intracellular Mlep outside vessels. 10μm bars.

**Supplemental Figure S2. Collision-induced dissociation of mycobacterial phenolic glycolipids (PGLs).** Related to Figures 3A and 3B.

(A) Collision-induced dissociation of an ammoniated adduct of PGL-1 standard isolated from *M. leprae* shows calculated masses on the structure (left) with detected masses shown in the chromatogram (right). 30V collision energy; blue diamond = collided ion. Detected ions are considered matches to calculated masses when they agree to within 10 parts per million (ppm) of mass accuracy.

(B) Collision-induced dissociation of PGL-*mar* standard isolated from wildtype *M. marinum*, shown as in A.

(C) Collision-induced dissociation of PGL-1 from total lipid extract of an *M. marinum:PGL-1* log phase culture, shown as in A. The fragment *m/z* 525, detected in collision of PGL-1 from *M. marinum:PGL-1* (C) and *M. leprae* (A), indicates the presence of a trisaccharide. Fragments detected at *m/z* 1140 in the PGL from *M. marinum:PGL-1* (C), and corresponding fragments at *m/z* 1574 and 1384 from *M. leprae* PGL-1 (A), indicate the sequential loss of sugars “c” and “b” from the lipid backbones.

**Supplemental Figure S3. Lower magnification image of nerve lesions.**

Related to Figure 3C. Lower magnification views of *mbp:eGFP-CAAX* larvae at 2 or 4 dpi with *M. leprae* and *M. marinum:PGL-1*, showing the apparently intact myelin sheath outside of the lesion (*M. marinum* shown for comparison).

A. 2 dpi, *M. leprae*
B. 4 dpi, *M. leprae*
C. 2 dpi, *M. marinum:PGL-1*
D. 2 dpi, wildtype *M. marinum*

**Supplemental Figure S4. Changes in axon ultrastructure in infected larvae.**

Related to Figure 4.

(A) Mean (±SEM) number of myelinated axons randomly selected (see Methods) in the hemi-spinal cords of larvae 2 days post-injection (dpi) with PBS control (white), *M. leprae* (Mlep; black) or *M. marinum:PGL-1* (Mm:PGL1; gray). 3 larvae per group.

(B) Mean (±SEM) number of total axons, quantified as in A.

(C) Two additional examples of myelin decompaction and dissociation in the spinal cords of Mlep-infected fish from Figure 4B. Highlights indicate myelinated axons (pink), nonmyelinated axons (orange), decompacted myelin (green highlight and arrows), and myelin dissociated from axons (blue highlight and arrows). 10μm bars.

**Supplemental Figure S5. Demyelinating lesion with a dead macrophage.**

Related to Figure 5I and Table S2. At 2 days post-infection, demyelinating lesions were identified in *M. marinum:PGL-1*-infected *mbp:eGFP-CAAX; mpeg1:Brainbow* larvae (Figure 5I). Each lesion was scored for presence or absence of infected macrophages (Table S2).

(A) Fish #7, which had 3 demyelinating lesions (dashed boxes).

(B) Insets of each lesion, showing myelin protrusions and colocalized, infected macrophages that are still intact (1 and 2) or fragmented (3) and dead. Arrowheads indicate myelin protrusions; for clarity, not every myelin protrusion that was scored is indicated with an arrowhead. White arrows, intact macrophages; yellow arrows, fragments of dsRed-positive macrophage membrane. 10μm bars.

**Supplemental Figure S6. Nitric oxide production in *M. marinum:PGL-1-*infected larvae**. Related to Figure 6.

(A) Representative confocal images of a macrophage aggregate in an *mpeg1* larva infected with *M. marinum:PGL-1* and stained with α-iNOS antibody. 10μm bar.

(B) Representative confocal images of a macrophage aggregate in larvae as in A, stained with α-nitrotyrosine antibody. 10μm bar.

## Movie Legends

**Movie S1. Rotating rendering of a demyelinating nerve lesion**. Related to Figures 2F and 2G. An *mbp:eGFP-CAAX* larva (green myelin) at 3 days post-fertilization was injected in the spinal cord with *M. leprae* (red), then imaged by confocal microscopy at 4 days post-infection. 10μm bar.

**Movie S2. Time lapse imaging of myelin protrusions forming from individual oligodendrocytes**. Related to Figure 3. Time-lapse movies of individual oligodendrocytes in the spinal cord of transient *mbp:eGFP-CAAX* transgenic larvae. Larvae were injected in the spinal cord near the labeled cell and imaged immediately for 12 hours. Injection sites are out of the frame to the left. 10μm bar; relative timecode.

(A) Injection of *M. marium:PGL-1*. Beginning ∼5 hours post-injection, the apparently intact myelin sheath bulges and 3 protrusions begin to form (arrows).

(B) Injection as in A of PBS vehicle.

**Movie S3. Timelapse confocal imaging of macrophages in the spinal cord of an uninjected larvae**. Related to Figure 5. An *mbp:eGFP-CAAX;mpeg1:Brainbow* larva (green myelin, red macrophages), at 4 days post-fertilization, imaged and displayed as in Movie S3. Myelin is shown as the fluorescence image and macrophages are shown as renderings. There is green autofluorescence from the skin dorsal to the myelinated axons, which covers the dorsal aorta. The autofluorescence is indicated by a rendered blue overlay. An arrow indicates a macrophage that associates with the myelin. Only a portion of the 12-hour movie is shown. 10μm bar; relative timecode.

**Movie S4. Timelapse confocal imaging of macrophages in the spinal cord of an uninjected larvae**. Related to Figure 5. Another example of an untouched larvae, imaged and displayed as in Movie S3. An arrow indicates a macrophage that associates with the myelin.

**Movie S5. Timelapse confocal imaging of macrophages in the spinal cord of a PBS-injected larvae**. Related to Figure 5. An *mbp:eGFP;mpeg1:Brainbow* larva (green myelin, red macrophages), at 4 days post-fertilization, was injected in the spinal cord with PBS + 2% phenol red and immediately imaged by confocal microscopy. Myelin is shown as the fluorescence image and macrophages are shown as renderings. Macrophages initially cluster at the injection site (circle) and crawl along myelinated axons (arrows); however, few or no macrophages remain associated with myelin after ∼8 hours post-injection. 10μm bar; relative timecode.

**Movie S6. Timelapse imaging of *M. leprae* injection in the spinal cord**.

Related to Figure 5C. An *mbp:eGFP-CAAX;mpeg1:Brainbow* larva (green myelin, red macrophages) at 4 days post-fertilization was injected in the spinal cord with *M. leprae* (cyan), imaged and displayed as in Movie S3. At the injection site (circle), a small aggregate of heavily-infected macrophages remain for the duration of imaging. Meanwhile, uninfected and infected macrophages, with bacilli visible inside of them, arrive at the injection site by crawling along or pushing apart myelinated axons (arrows). After phagocytosing bacilli or contacting other cells, infected macrophages crawl away from the injection site, carrying *M. leprae*. Myelin is shown as the fluorescence image; macrophages are shown as transparent renderings to show the rendered *M. leprae* inside. The imaged area, centered around the injection site, has dimensions x = 200μm, y = 95μm, z = 50μm. 10μm bar; relative timecode.

**Movie S7. Timelapse imaging of *M. marinum* injection in the spinal cord**.

Related to Figure 5D through 5F. An *mbp:eGFP-CAAX;mpeg1:Brainbow* larva (green myelin, red macrophages cells), at 4 days post-fertilization, was injected with ∼100 colony-forming units of *M. marinum* (Mm, cyan), imaged and displayed as in Movie S3. The macrophage response is similar to that in *M. leprae* infection, with several heavily-infected macrophages remaining at the injection site (circle) for most of imaging. Macrophages crawl along the myelin sheaths (arrows) toward the injection site, phagocytose bacilli, and exit the frame. Macrophage morphology, speed, and myelin colocalization are quantified in Figures 5D through 5F.

**Movie S8. Timelapse imaging of *M. marinum:PGL-1* injection in the spinal cord**. Related to Figures 5D through 5F. An *mbp:eGFP-CAAX;mpeg1:Brainbow* larva (green myelin, red macrophages), at 4 days post-fertilization, was injected with ∼100 colony-forming units of *M. marinum:PGL-1* (Mm:PGL1; cyan), imaged and displayed as in Movie S3. As with *M. leprae* and Mm injection, several heavily-infected macrophages remain at the injection site (circle). Macrophages crawl along the myelin sheaths (arrows) toward the injection site, phagocytose bacilli, and exit the frame for distal sites. Macrophage morphology, speed, and myelin colocalization appear similar to that in *M. marinum* infection (quantified in Figures 5D through 5F).

## STAR Methods

### Zebrafish husbandry, infections, and drug treatment

Zebrafish husbandry and experiments were conducted in compliance with guidelines from the U.S. National Institutes of Health and approved by the University of Washington Institutional Animal Care and Use Committee, the Office of Animal Research Oversight of the University of California Los Angeles, and the Institutional Biosafety Committee of the University of California Los Angeles. Wildtype AB strain larvae or transgenics in the AB background were used, including Tg(*kdrl:dsRed*)^s843^ (Jin et al., 2005), Tg(*mbp:CAAX-GFP*)^ue2Tg^ (Almeida et al., 2011), Tg(*mpeg1:Brainbow*)^w201^ (Pagán et al., 2015), Tg(*lysC:EGFP*)^nz117^ (Hall et al., 2007) and Tg(*mpeg1:YFP*)^w200^ (Roca and Ramakrishnan, 2013). Larvae were anesthetized with tricaine (MS-222, Sigma) as described (Takaki et al., 2013), prior to imaging or infection. Larvae were infected via the caudal vein or hindbrain ventricle at 2 dpf as previously described (Takaki et al., 2013), or infected in the ventral spinal cord adjacent to the cloaca at 2-4 dpf. Nitrotyrosine and iNOS were detected in infected larvae as described (Cambier et al., 2014; Elks et al., 2014). Larvae were treated with nitric oxide mediators by soaking in fish water beginning 4-6 hours after infection to allow macrophage recruitment to the injection site; drugs were re-added every 12 hours. L-NAME (1000uM) and cPTIO (500uM) were used to inhibit iNOS and scavenge reactive oxygen/nitrogen species, as described (Cambier et al., 2014). SNAP (100uM) and spermine NONOate (10uM) were used to exogenously add nitric oxide, as described (Kong et al., 2016; Siamwala et al., 2012).

### Cell culture

BMDMs were isolated and differentiated from C57Bl/6 mice. Briefly, bone marrow isolated from femurs and tibias from male mice at 6-10 weeks of age were cultured in media consisting of DMEM with 20% FBS, CMG conditioned media containing M-CSF, and 1X Pen/Strep for 6 days. All media was washed twice with PBS, then antibiotic-free BMDM media was added prior to addition of bacteria. Cells were infected with Mlep (harvested from footpads of nude mice) at indicated MOI or with equivalent volumes of log-phase (OD_600_ ≈ 0.5) wildtype *M. marinum* or *M. marinum:PGL-1* cultures (growth conditions described below). MOI was obtained by growing the cultures on 7H10 plates. Cells were harvested and RNA isolated as described (Teles, 2013).

### Morpholino, RNA, and liposome injections

Morpholinos (Gene Tools) were used to block translation or splicing of transcript for *irf8* (0.6mM of sequence AATGTTTCGCTTACTTTGAAAATGG) (Li et al., 2011), *pu.1* (mixture of 0.375mM of sequence CCTCCATTCTGTACGGATGCAGCAT and 0.025mM of sequence GGTCTTTCTCCTTACCATGCTCTCC) (Clay et al., 2007), *MyD88* (5mM of sequence GTTAAACACTGACCCTGTGGATCAT) (Bates et al., 2007; Cambier et al., 2014), or *CCR2* (0.3mM of sequence AACTACTGTTTTGTGTCGCCGAC) (Cambier et al., 2014). Morpholinos or *in* vitro-transcribed H2B-CFP (Megason, 2009) were diluted in a 1x Buffer Tango (Thermo Scientific) containing 2% phenol red (Sigma) and injected into the yolk of 1-2 cell-stage embryos in ∼1 nL (Tobin et al., 2012). Lipo-PBS and lipo-clodronate (http://clodronateliposomes.org) (van Rooijen et al., 1996) were diluted in PBS + 2% phenol red and injected into 2-dpf-old larvae in ∼10 nL via the caudal vein. To generate larvae with fluorescent mitochondria in axons, eggs were coinjected with 50ug/uL *tol2* transposase RNA, 25ng/uL of an existing pDEST-UAS:MLS-dsRed plasmid (O’Donnell et al., 2013), and 25ng/uL of a constructed pDEST plasmid consisting of GAL4 expressed from the *Xenopus laevis* neuronal beta tubulin (*nbt)* promoter (Peri and Nüsslein-Volhard, 2008). To generate larvae with individual labeled oligodendrocytes, eggs were coinjected with *tol2* RNA and 1ng/uL of the mbp:eGFP-CAAX plasmid (Almeida et al., 2011).

### Bacterial Strains

*M. marinum* M strain (ATCC #BAA-535) and its derivative, *M. marinum:PGL-1*, expressing tdTomato, wasabi or eBFP under control of the msp12 promoter (Cosma et al., 2005; Takaki et al., 2013), were grown under hygromycin (Mediatech) or kanamycin (Sigma) selection in 7H9 Middlebrook medium (Difco) supplemented with oleic acid, albumin, dextrose, and Tween-80 (Sigma) (Takaki et al., 2013). *M. marinum:PGL-1* was constructed by transforming *M. marinum* with the integrating plasmid pWM122, which encodes the *M. leprae* genes ML0126, ML0127, ML0128, ML2346c, ML2347, and ML2348 under the *M. fortuitum pBlaF\** promoter (Tabouret et al., 2010). Kanamycin-resistant transformants were confirmed by PCR using primers targeting all six *M. leprae* genes (Tabouret et al., 2010). A single transformant was further confirmed by mass spectrometry of its phenolic glycolipids; this strain was used for all zebrafish experiments. For infections, *M. leprae* was isolated from mouse footpads, labeled with fluorescent dye (PKH67, PKH29, or CellVue Claret, Sigma), then tested for viability by radiorespirometry, as described (Lahiri et al., 2005). *P. aeruginosa* PAO1 expressing GFP has been described (Brannon et al., 2009).

### Mass spectrometry

*M. marinum* wildtype and *M. marinum:PGL-1* were cultured in 20 mL of 7H9 medium, supplemented with 10% albumin/dextrose/catalase (EMD Chemicals, San Diego, CA) to mid-log phase (OD_600_ = 0.5). Total lipids were extracted from cell pellets as described (Layre et al., 2011) using LC-MS grade solvents (Fisher) and dried under nitrogen. Each lipid extract, in addition to PGL-1 standard from *M. leprae* (BEI), was analyzed on an Agilent Technologies 6520 Accurate-Mass Q-Tof and a 1200 series HPLC system with a Varian Monochrom diol column (3 **μ**m × 150 mm × 2 mm) and a Varian Monochrom diol guard column (3 **μ**m × 4.6 mm). Lipids were resuspended at 0.5 **μ**g/mL in solvent A (hexanes:isopropanol, 70:30 [v: v], 0.02% [m/v] formic acid, 0.01% [m/v] ammonium hydroxide), then 10 **μ**g were injected and the column was eluted at 0.15 mL/min with a binary gradient from 0% to 100% solvent B (isopropanol:methanol, 70:30 [v/v], 0.02% [m/v] formic acid, 0.01% [m/v] ammonium hydroxide): 0–10 min, 0% B; 17–22 min, 50% B; 30–35 min, 100% B; 40–44 min, 0% B, followed by additional 6 min 0% B postrun. Ionization was maintained at 325° C with a 5 L/min drying gas flow, a 30 psig nebulizer pressure, and 5,500 V. Spectra were collected in positive ion mode from *m/z* 100 to 3,000 at 1 spectrum/s. Continuous infusion calibrants included *m/z* 121.050873 and 922.009798 in positive ion mode. Collision-induced dissociation was performed with an energy of 30 V.

### Fluorescence microscopy

Wide-field microscopy was performed using a Nikon Eclipse Ti-E equipped with a C-HGFIE 130W mercury light source, Chroma FITC (41001) filter, and ×2/0.10 Plan Apochromat objective. Fluorescence images for evaluating bacterial escape from the vasculature were captured with a CoolSNAP HQ2 Monochrome Camera (Photometrics) using NIS-Elements (version 3.22). Quantification of fluorescent bacterial infection, using Fluorescent Pixel Count (FPC) quantification of images of individual embryos, was performed as previously described (Takaki et al., 2012).

### Confocal microscopy

For confocal imaging, larvae were imbedded in 1.5% low melting-point agarose (Davis and Ramakrishnan, 2009). A series of z-stack images with a 2-3 μm step size was generated through the infected spinal cord with the image centered at the injection site or cloaca, using either the galvo scanner (laser scanner) of the Nikon A1 confocal microscope with a ×20 Plan Apo 0.75 NA objective, or the resonant laser scanner of a Leica TCS-SP5 AOBS confocal microscope with a 20x Plan Apo 0.70 NA. Bacterial burdens were determined by using the three-dimensional surface-rendering feature of Imaris (Bitplane Scientific Software) (Yang et al., 2012). Macrophage numbers, shape and speed were determined using tracking of surface-rendered features on Imaris. When events were compared between larvae, identical confocal laser settings, software settings and Imaris surface-rendering algorithims were used.

### Transmission electron microscopy

Before preparing larvae for TEM, they were imaged by confocal microscopy in order measure the distance from the cloaca to the spinal cord lesion; this allowed sections to be taken through confirmed demyelinating lesions after the larvae were fixed. After rescuing larvae from 1.5% agarose used for confocal imaging, healthy larvae were anesthetized, cooled to 4°C, then fixed overnight in ice-cold 0.1 M sodium cacodylate containing 2% glutaraldehyde, 4% paraformaldehyde and 4% sucrose (Czopka and Lyons, 2011). Following several washes in buffer, the larvae were postfixed in a solution of 2% osmium tetroxide and 0.1M imidazole in buffer for 1 hour on ice. The larvae were rinsed multiple times in water and treated with 0.5% uranyl acetate overnight at 4^°^C. They were then dehydrated through a graded series of ethanols, passed through propylene oxide and infiltrated with Eponate12 (Ted Pella) overnight. The larvae were embedded in fresh Eponate12 and the blocks polymerized at 60^°^C. The areas of interest were identified relative to the cloaca, and 50nm (silver interference color) sections were taken through these areas on an ultramictome (RMC MTX) and deposited on grids. The grids were stained with saturated uranyl acetate and Reynolds lead citrate and examined on a JEOL 100CX electron microscope at 60kV. Images were collected on film, and then scanned at 1200 dpi to create digital files. Axons were identified by the presence of microtubules and/or microfilaments and an intact outer membrane. Decompacted myelin was identified by the presence of large, electron-lucent spaces in between myelin lamellae that were not observed in the absence of infection. Myelin dissociated from axons was identified by the presence of electron dense “whorls” of myelin lamellae that did not contain an axon. Mitochondria were identified by an intact double membrane and cristae. Axon number, myelination, size, and presence of mitochondria were scored by randomly selecting axons in each image. To assure that axons selection was random, each image was opened in its original dimensions in Adobe PhotoShop and overlaid with a 50-pixel grid; only axons under grid nodes were scored (Czopka and Lyons, 2011).

### Statistics

Statistical analyses were performed on Prism (version 5.0a, GraphPad). Not significant, p ≥ 0.05; * p < 0.05; ** p < 0.01; *** p < 0.001; **** p < 0.0001.

